# Plasticity as an Underlying Mechanism of Tumor Heterogeneity in Breast Cancer

**DOI:** 10.1101/2020.03.11.987511

**Authors:** Muhammad Waqas Akbar, Murat Isbilen, Baris Kucukkaraduman, Secil Demirkol Canli, Ege Dedeoglu, Shila Azizolli, Isli Cela, Abbas Guven Akcay, Hasim Hakanoglu, Ali Osmay Gure

**Affiliations:** Department of Molecular Biology and Genetics, Bilkent University, Ankara, Turkey; DNAFect Genetics Consulting R&D and Biotechnology Inc., Kocaeli, Turkey; Molecular Pathology Application and Research Center, Hacettepe University, Ankara, Turkey

## Abstract

Breast cancer shows plasticity in terms of classification. Upon drug treatment and metastasis some tumors switch to another subtype leading to loss of response to therapy. In this study, we ask the question which molecular subclasses of breast cancer are more switchable upon drug therapy and metastasis. We used in silico data to classify breast cancer tumors in PAM50 molecular classes before treatment and after treatment using gene expression data. Similar analysis was performed for primary tumors and their metastatic growth. Our analysis showed that in both scenarios some breast tumors shift from one class to another. This suggests that patients who underwent chemotherapy but resulted in relapse or/and metastasis should be retyped for molecular subclass so that treatment protocol should be adopted according to those subtypes. Additionally, 20 genes were identified as biomarkers for metastasis in breast cancer.

## Introduction

Breast cancer (BC) is the second leading cause for cancer related mortality in women after lung cancer^1,2^. In clinics, breast cancer is classified using estrogen receptor (ER), progesterone receptor (PGR) and Erb-B2 Receptor Tyrosine Kinase 2 (ERBB2, previously known as HER2) statuses through immunohistochemistry (IHC classes). Moreover, BC is divided in Luminal A, Luminal B, HER2 enriched, Basal and Normal like subgroups by molecular classification ^3^. Metastatic breast cancer (MBC) is the type of BC in which cancer cells leave primary cancer tissue and spread to other parts of the body. MBC remains an incurable disease as its median overall survival is about 3 years and 25% patients show only 5 year survival^4^.

MBC consists of a series of sequential steps, including transfer of cells from primary tumor to circulation (intravasation), survival in circulation, extravasation in a new organ, colonization and metastatic growth. Metastatic cells go through epithelial to mesenchymal transition (EMT) to achieve metastasis^5^. This metastatic cascade results in switching the IHC status of BC in some samples, assigning metastatic growth a different class for which primary therapy might not be suitable and a different protocol should be opted to combat this growth^6^. Previously similar subtype switching is observed in PAM50 classes as well ^7^. In this paper we address this switching as plasticity. Additionally, this subtype plasticity is also seen among pre- and post-treated tumors due to epithelial to mesenchymal transition^8^.

In this paper we used public gene expression datasets to determine if there is a subtype switch between pre- and post-therapy patient tumor samples and also if this switch is present between primary and metastatic tissue. Moreover MBC biomarkers were identified using primary tumor expression data which can be used to determine if a breast cancer is capable of metastasis or not.

## Methods

### PAM50 plasticity

Datasets GSE10281, GSE32518, GSE32072 (patient samples before and after treatment), GSE32518 (patient samples collected with fine needle aspiration and core needle biopsy), GSE125989, GSE110590 (patient samples for primary and metastatic tissues) were downloaded from GEO NCBI database. All datasets were “justRMA” normalized using BRB Array Tools^9^. PAM50 signature was used to classify breast cancer into 5 molecular subgroups: Luminal A (LumA), Luminal B (LumB), HER2 enriched (HER2), Basal and Normal like (Normal) as described previously^10^ for each dataset. Paired samples were matched for pre- and post-therapy, with fine needle aspiration and core needle biopsy, or primary and metastatic tissue to compare PAM50 plasticity in these samples.

### Metastasis biomarkers

For metastasis-free survival analysis, GSE2603 (n=82, treatment not disclosed), GSE6532 ((n= 293, platform: Affymetrix Human Genome U133A and B Array, Tamoxifen treated), (n=87, Affymetrix Human Genome U133 Plus 2.0 Array)), GSE7390 (n=198, systemically untreated patients), GSE11121 (n=200, only surgery was performed), GSE12276 (n=204, adjuvant chemotherapy treated), GSE20685 (n=327, adjuvant chemotherapy treated) and GSE58812 (n=108, adjuvant chemotherapy treated) were downloaded. All datasets were “justRMA” normalized using BRB Array Tools. We performed cox proportional hazard analysis using “Survival” package built in R software for all datasets^11,12^. Genes that showed same direction in all datasets and were significant for 5/8 analysis were selected as metastasis biomarkers. 40 probesets were found significant and to select top 10 good and bad prognostic genes, genes were ranked by coxp value in each dataset individually for both directions and these ranks were summed. Top 10 genes were then selected from good prognostic genes based upon the least ranksum score and another 10 genes were selected from bad prognostic genes based on the same criteria. For genes which were significant with 2 or more probesets, only one probeset with the least ranksum score was selected.

## Results

### PAM50 plasticity

PAM50 is also prone to show different classes, as analyzed in dataset GSE32518, where authors collected fine needle aspiration and core needle biopsies from matching patients (N=37). When these samples were typed for PAM50, 78% of these samples (n=29) showed same phenotype in both biopsy methods while 22% samples (n=8) showed discordant subtypes. One HER2 sample by core needle biopsy was identified as Basal and another one was identified as LumB. One sample was identified as LumA (Core needle biopsy) and LumB (Fine needle aspiration). Most discordant results were for Normal subtype (Core needle biopsy) where 4 samples were identified Basal and 1 sample as LumA (Table 1). This PAM50 discordance can be attributed to tumor heterogeneity.

**Table 1:** Comparison of PAM50 subtyping for samples collected through fine needle aspiration and core needle biopsy (GSE32518).

|  |  | Fine needle aspiration |  |  |  |  |
| --- | --- | --- | --- | --- | --- | --- |
|  |  | Basal | HER2 | LumA | LumB | Normal |
| Core needle biopsies | Basal | 7 | 0 | 0 | 0 | 0 |
|  | HER2 | 1 | 5 | 0 | 1 | 0 |
|  | LumA | 0 | 0 | 9 | 1 | 0 |
|  | LumB | 0 | 0 | 0 | 7 | 0 |
|  | Normal | 4 | 0 | 1 | 0 | 1 |

Plasticity exists between PAM50 classes, and tumor subtype can switch to another upon drug treatment through EMT and immunological status. To study PAM50 plasticity, we analyzed different datasets in which matched tumor samples gene expression data was available for both pre- and post-chemotherapy. When Letrozole pre- and post-treated samples (GSE10281) were compared for PAM50 class plasticity, 39% samples (7/18) retained their PAM50 class before and after Letrozole therapy and 61% (11/18) samples changed their class. Basal (n=2) and LumB (n=1) subtype was most stable in this dataset. One HER2, 2 LumA and 1 normal subtype samples retained their subtype while 3 LumA, 3 HER2 and 4 normal pre therapy samples switched to another subtype. Most of these samples (n=6) switched to LumB and 4 samples switched to Basal subtype (Table 2). Authors reported in their paper that upon chemotherapy treatment, residual tumors attain mesenchymal properties as expected^13^. Among patients treated with Anthracyclines and Taxanes based neoadjuvant chemotherapy (GSE32518), 2 basal samples, 1 HER2, 3 LumA, 3 lumB and 1 Normal retained their subtype and 2 basal, 3 HER2, 4 LumA, 8 LumB and 1 normal switched their phenotype (Table 3). Most samples switched their subtype to normal like tumors. Authors reported that genes related to extracellular metabolism, angiogenesis and developmental processes are related to chemoresistance in these patients^14^. In another dataset (GSE32072), where gene expression data for 21 cancer samples was available for pre- and post-neoadjuvant therapy (9 patients anthracycline based therapy, 8 patients anthracycline/taxane based therapy and 2 patients trastuzumab therapy). Additionally, 4 patients were also included which were not treated with any neoadjuvant therapy as control. Upon PAM50 subtyping, patients who did not receive any neoadjuvant chemotherapy, retained their classes (2 basal and 2 LumA). Interestingly, all basal (n=5), Her2 (n=4) and LumB (1) retained their subtype. Out of 6 pre-therapy lumA samples only one changed to LumB subtype. Normal subtype samples changed the most to other subtypes in this dataset (N=4) and only one sample retained its phenotype (Table 4). Authors reported enrichment of energy and metabolism related genes in residual samples while immune related genes were depleted^15^.

**Table 2:**
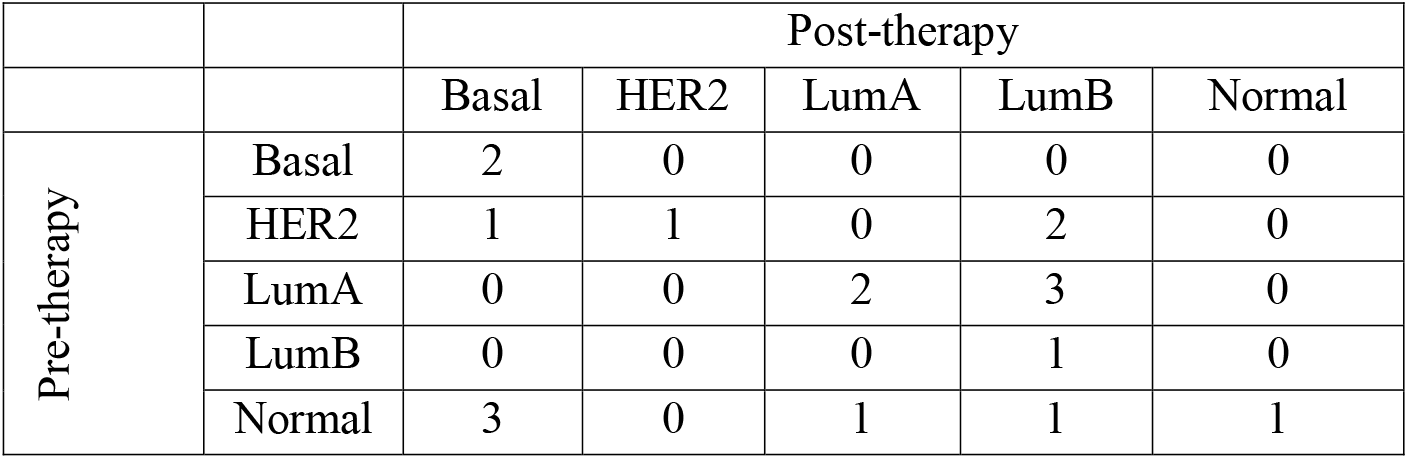
Comparison of PAM50 subtypes pre- and post-Letrozole therapy (GSE10281).

|  |  | Post-therapy |  |  |  |  |
| --- | --- | --- | --- | --- | --- | --- |
|  |  | Basal | HER2 | LumA | LumB | Normal |
| Pre-therapy | Basal | 2 | 0 | 0 | 0 | 0 |
|  | HER2 | 1 | 1 | 0 | 2 | 0 |
|  | LumA | 0 | 0 | 2 | 3 | 0 |
|  | LumB | 0 | 0 | 0 | 1 | 0 |
|  | Normal | 3 | 0 | 1 | 1 | 1 |

**Table 3:** Comparison of PAM50 subtypes pre- and post-Anthracyclines and Taxanes based neoadjuvant chemotherapy (GSE32518).

|  |  | Post-therapy |  |  |  |  |
| --- | --- | --- | --- | --- | --- | --- |
|  |  | Basal | HER2 | LumA | LumB | Normal |
| Pre-therapy | Basal | 2 | 0 | 0 | 0 | 2 |
|  | HER2 | 1 | 1 | 0 | 1 | 1 |
|  | LumA | 0 | 0 | 3 | 0 | 4 |
|  | LumB | 1 | 0 | 4 | 3 | 3 |
|  | Normal | 0 | 0 | 1 | 0 | 1 |

**Table 4:** Comparison of PAM50 subtypes pre- and post-neoadjuvant chemotherapy (GSE32072).

|  |  | Post-therapy |  |  |  |  |
| --- | --- | --- | --- | --- | --- | --- |
|  |  | Basal | HER2 | LumA | LumB | Normal |
| Pre-therapy | Basal | 5 | 0 | 0 | 0 | 0 |
|  | HER2 | 0 | 4 | 0 | 0 | 0 |
|  | LumA | 0 | 0 | 5 | 1 | 0 |
|  | LumB | 0 | 0 | 0 | 1 | 0 |
|  | Normal | 2 | 1 | 1 | 0 | 1 |

Breast cancer metastasis can be molecularly distinct from their primary tissues^16^. Metastatic samples can switch to another PAM50 class when compared with primary tissue class. In dataset GSE125989, 16 matched tissues were available for both brain metastasis and primary site. These samples were PAM50 subtyped. 50% (8) of them retained their phenotype when primary site vs metastasis PAM50 classes were compared. One Basal changed to Her2 and 1 Her2 changed to basal. Major subtype switching was observed in Normal primary tissue phenotype, where 3 metastasized tissues became basal and 2 became HER2 (Table 5). Authors showed that when gene expression of metastasized tissue was compared with primary tissue, mesenchymal markers were up regulated while epithelial markers were downregulated. Additionally tumor infiltrating lymphocytes, B cells and dendritic cell gene signatures were downregulated in metastatic tissues. Dataset GSE110590 contains gene expression for 16 primary breast cancer tissues and their 67 matched metastasis. 55% (37/67) metastatic tissues retained the same PAM50 class when compared with primary site. Most stable classes were basal where 85% (23/27) metastatic sites retained basal phenotype and HER2 100% (4/4). While LumA showed 32% (10/31) and Normal showed 0% (0/5) consistecy. Most primary samples when metastasized switched to Basal and LumB class (Table 6).

**Table 5:** Comparison of PAM50 subtyping for primary tissues vs. brain metastatic tissues (GSE125989).

|  |  | Metastasized tissue |  |  |  |  |
| --- | --- | --- | --- | --- | --- | --- |
|  |  | Basal | HER2 | LumA | LumB | Normal |
| Primary tissue | Basal | 2 | 1 | 0 | 0 | 0 |
|  | HER2 | 1 | 2 | 0 | 0 | 1 |
|  | LumA | 0 | 0 | 4 | 0 | 0 |
|  | LumB | 0 | 0 | 0 | 0 | 0 |
|  | Normal | 3 | 2 | 0 | 0 | 0 |

**Table 6:** Comparison of PAM50 subtyping for primary tissues vs. metastatic tissues (GSE110590).

|  |  | PAM50 metastasis |  |  |  |  |
| --- | --- | --- | --- | --- | --- | --- |
| Sample name | Primary tissue | Basal | HER2 | LumA | LumB | Normal |
| A11 | Basal | 5 | 0 | 0 | 0 | 0 |
| A15 | Basal | 2 | 0 | 0 | 0 | 3 |
| A1 | Basal | 4 | 0 | 0 | 0 | 1 |
| A23 | Basal | 5 | 0 | 0 | 0 | 0 |
| A5 | Basal | 2 | 0 | 0 | 0 | 0 |
| A7 | Basal | 5 | 0 | 0 | 0 | 0 |
| A8 | HER2 | 0 | 4 | 0 | 0 | 0 |
| A12 | LumA | 0 | 0 | 2 | 4 | 0 |
| A17 | LumA | 2 | 0 | 0 | 0 | 0 |
| A26 | LumA | 3 | 0 | 0 | 0 | 0 |
| A28 | LumA | 0 | 0 | 3 | 3 | 0 |
| A2 | LumA | 0 | 1 | 0 | 2 | 0 |
| A30 | LumA | 5 | 0 | 0 | 0 | 0 |
| A34 | LumA | 0 | 0 | 4 | 0 | 0 |
| A4 | LumA | 0 | 1 | 1 | 0 | 0 |
| A20 | Normal | 5 | 0 | 0 | 0 | 0 |

In another study, authors compared 123 patients’ primary tissues with metastatic tissues for PAM50 switching. Metastatic tissues included skin, lymph node, liver, bone, lung, peritoneum and others. Subtype concordance for Basal was highest (100%), followed by HER2 (76.92%) and LumB (70%). The most switchable subtype was LumA where 55.32% sample switched to luminal B and HER2^7^.

Above analysis suggests PAM50 subtype switch between different classes both due to chemotherapy treatment and metastasis. Therfore subtyping for residual cancer or metastasis should be re-evaluated. Additionally, basal phenotype is the most stable phenotype and LumA is the most unstable one. Moreover PAM50 classification is subjected to sampling method as we show that different sampling techniques can result in different classification of the same samples.

### Biomarkers for metastasis

To identify biomarkers for metastasis, we used 7 cohorts for which metastasis-free survival data was reported. Treatment protocol and patient numbers for each cohort are described in methods section. We selected 10 good and 10 bad prognostic genes. All selected 20 genes and their respective probeset IDs are shown in figure 1.

**Figure 1:**
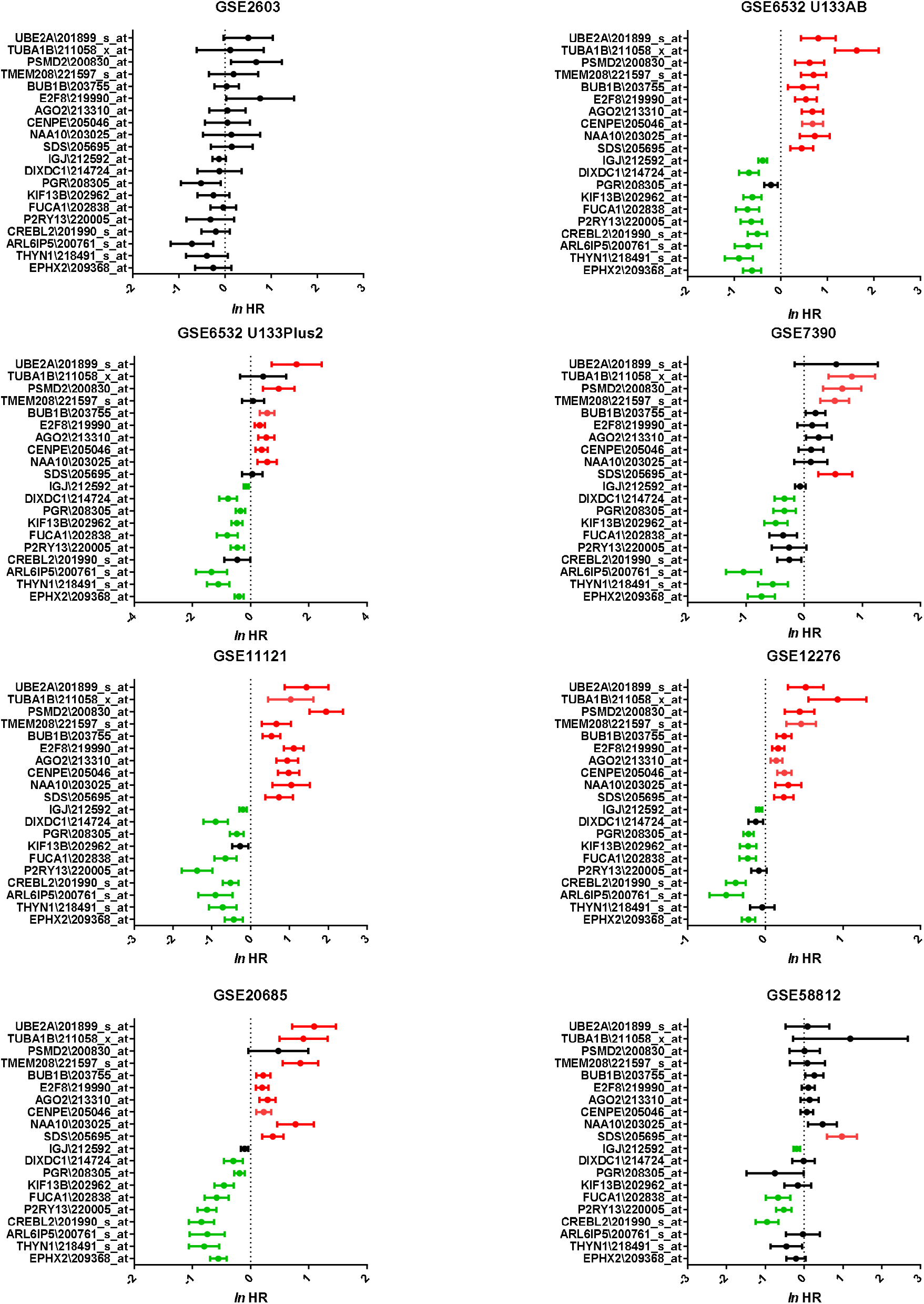
Metastasis-free survival analysis: 20 genes (10 good prognostic, 10 bad prognostic) were selected as metastasis biomarkers. Gene names and probe IDs are shown on y axis. Y axis shows Hazard ratio (HR) in natural logarithm with 95% CI. Genes which are good prognostic and significant are shown in as green while bad prognostic and significant genes are shown in red color. If a gene was not significant it is shown in black.

Our selected genes can predict metastasis-free survival or distant metastasis-free survival in several cohorts. Next we used these genes to cluster patients in metastatic prone or resistant groups with the aim to identify enriched gene sets among these using GSEA^17^. Surprisingly, no common gene sets enriched in metastatic prone and resistant groups were found, suggesting that different mechanisms like EMT, inflammation, cell cycle and immune response are involved in metastasis (data not shown).

### Good prognostic genes

Among the 20 genes selected, 10 were good prognostic. Epoxide hydrolase 2 (EPHX2) is largely involved in metabolism of epoxy fatty acids and plays an important part in cardiovascular biology, renal function, inflammation response and pain^18^. Previously, EPHX2 has been associated with metastasis suppression in breast cancer and its expression was four-fold higher in lymph node negative tumors as compared with lymph node positive tumors^19^. In another meta-analysis for breast cancer metastasis, EPHX2 was identified as metastasis suppressor gene and its biological functions were associated with drug metabolism, inflammatory response and reactive oxygen species metabolism^20^. So, it might be possible that EPHX2’s diverse role in metabolism of carcinogens might play a role in metastasis of breast cancer. Previously high glucose levels were associated with increased breast cancer proliferation and metastasis^21^ and interestingly specificity protein 1 (sp1) is shown to downregulate EPHX2 under increased glucose levels^22^. EPHX2 role has also been associated with recurrence in breast cancer^23^.

Thymocyte nuclear protein 1 (THYN1), also known as THY28, is a highly conserved protein from bacteria to mammals and plays an important role in B cell development^24^ and its downregulation has been associated with induction of apoptosis in Ramos B lymphoma cells^25^. However its role in breast cancer is not well studied. Previously, another study on breast cancer metastasis has associated its role as good prognosis marker as they used the same cohort GSE6532 (also used in this study). In another study, TYN1 role was associated with recurrence in breast cancer^23^. THYN1 is also negatively correlated with numerical chromosomal heterogeneity in NCI60 cancer cell lines. In this study, authors showed that chromosomal instability is associated with EMT, cancer invasiveness and metastasis^26^.

ADP ribosylation factor like GTPase 6 interacting protein 5 (ARL6IP5), also known as JWA, has been shown previously as microtubule associated protein and has been related to differentiation, apoptosis and response to stress stimulators and its expression is mediated by ER^27^. In NIH3T3 and HELF cell lines, increased H_2_O_2_ levels resulted in increased expression of ARL6IP5 expression indicating its role in oxidative stress^28^. In breast cancer, knockdown of ARL6IP5, resulted in increased invasion and migration along with decreased apoptosis in MDA-MB-231 cells^29^. In another study, authors showed that expression of CXCR4 (breast cancer metastatic marker) was inversely proportional to ARL6IP5 and ARL6IP5 decreases breast cancer migration and invasion by proteomic degradation of CXCR4^30^. Moreover, ARL6IP5 has been shown to suppress metastasis of melanoma cells as well as cancer cell adhesion, invasion and metastasis in gastric cells, similar to our analysis^31–33^.

CAMP responsive element binding protein like 2 (CREBL2) is identified as a metabolic regulator^34^. CREBL2 was shown to be expressed by T47D cell line when compared with MDA-MB-435 (a highly metastatic cell line)^35^. CREBL2 expression is decreased in inflammatory breast cancer (an aggressive subtype of breast cancer)^36^. CREBL2 was found to be downregulated in OVCAR paclitaxel resistant cell line (ovarian cancer cell line) when compared with controls^37^. CREBL2 was also found downregulated among grade 3 breast cancer samples^38^. In another study, to identify genes differentially expressed between patients who suffered from distant metastasis and ones who did not, CREBL2 was downregulated in metastatic group^39^.

Purinergic receptor P2Y13 (P2RY13) is a G-protein-coupled receptor and it is downregulated upon EGF- and hypoxia-induced EMT in breast cancer cells^40,41^. Additionally, overexpression of P2RY13 is associated with better recurrence free survival in hepatocellular carcinoma^42^.

Alpha-L-Fucosidase 1(FUCA1) is a lysosomal enzyme involved in cleavage of terminal fructose residue. Its lower expression has been associated with aggressive tumors such as breast, neuroblastoma and colorectal cancer. Moreover its reduced expression is related to shorter recurrence and cancer free survival in lymph node positive breast cancer luminal B patients^43^. Its lower expression is also related to metastasis in solid tumors^44^. One study showed that FUCA1 functions downstream of p53 and can contribute to EGFR signaling repression by removing fucose from EGFR and thus is a marker of lower survival in breast cancer^45^. Among triple negative breast cancer patients, FUCA1 downregulation is associated with decreased survival through modulation of glycosylation status^46^.

Kinesin family member 13B (KIF13B) belongs to Kinesin 3 family. KIF13B is downregulated upon estrogen stimulation of breast cancer cell proliferation^47^. And its expression is upregulated in paclitaxel resistant MCF7 cells when compared with control cells^48^. Although KIF13B is not studied well in cancer, its role was emphasized as VEGFR2 trafficker in endothelial cells previously. Thus it plays an important role in angiogenesis and its depletion leads to reduced VEGF induced capillary tube formation, endothelial cells migration and neovascularization in mice^49^.

Progesterone receptor (PGR) status has been described in previous sections as it goes through plasticity during metastasis. In one study, authors found 46% tumors shifting from PR positive status to negative status in metastatic tissue, although in the same study they also saw the opposite trend for 16% tumors as well^50^. In another study, 40.4% discordance was found between primary and metastatic tissue regarding PGR status and 74% discordance was due to loss of PGR status^51^.

DIX domain containing 1 (DIXDC1) gene is a regulator of Wnt signaling pathway and its expression has been associated with good prognosis in breast cancer as well as metastasis repression in lung adenocarcinoma as expected^52,53^. Knockdown of DIXDC1 in glioma cells has been shown to inhibit proliferation and migration of these cells^54^. Interestingly lower expression of DIXDC1 has been associated with worse outcomes in hepatocellular carcinoma^55^ and intestinal type gastric carcinoma^56^.

Joining chain of multimeric IgA and IgM (JCHAIN), previous symbol IGJ, is downregulated in ductal carcinoma *in situ* and invasive ductal carcinoma when compared with normal stroma^57^. Additionally, JCHAIN was previously shown to be downregulated in both tumor stroma and tumor epithelium in inflammatory breast cancer (aggressive form of breast cancer related to metastasis) when compared with non-inflammatory breast cancer^36^.

### Bad prognostic genes

Serine dehydratase (SDS) converts L-serine to pyruvate and ammonia. It has not been studied extensively in breast cancer. In one study, its expression was shown to be higher in tumor samples when compared with normal samples^58^.

N-Alpha-Acetyltransferase 10, NatA Catalytic Subunit (NAA10), also known as ARD1, is related to post-translational modifications and has been previously shown to increase breast cancer proliferation upon overexpression^59^. Its expression is increased in breast tumor cells when compared with normal adjacent cells^60^. In one study, authors related its expression with lymph node metastasis in breast cancer^60^. Surprisingly in another study, it was reported as the inhibitor of metastasis by inhibiting cell motility^61^.

Centromere protein E (CENPE), with chromosome misalignment being the hallmark of cancer, aids the proliferation of tumor cells via its involvement in capturing and positioning the chromosomes to mitotic spindle^62,63^. The inhibition of CENP-E causes the death and remediation of triple negative breast cancer cells^64,65^.

Argonaute-2 (AGO2) plays an indispensable role in the post-transcriptional downregulation of gene expression through guided small-RNAs. It is one of the four members of the Argonaute protein family which is highly conserved amongst different species and is the only member ofto have catalytic activity and therefore, is the most important for gene silencing along with the RISC complex^66,67^. AGO2 has been linked to different types of cancers previously^68,69^. AGO2 expression was shown to increase in Her2 and basal subtype. Additionally increase in AGO2 expression showed reduced relapse free survival. In the same research it was indicated that cancer cell-lines showed weak AGO2 expression as opposed to the tissue samples, suggesting a role of the tumor microenvironment in AGO2 expression ^70^. AGO2 has also been shown to be highly expressed in a subset of ERα+ breast tumors and is correlated with poor prognosis in patients. It was shown that the endocrine resistance of ER+/PGR-tumors were highly correlated with AGO2 overexpression^71^. Cells transfected with AGO2 were shown to have a more aggressive phenotype with higher proliferative capabilities and higher expression of pro-oncogenic microRNAs^72^.

E2F8 is the eighth member of the E2F transcription factors, which are expressed in a variety of tissues and is involved in regulatory functions of cellular proliferation, differentiation, cell apoptosis, DNA repair & cell cycle ^73,74^. E2F8 plays a major role in cell-cycle progression, especially to the G1 to S phase transition by binding to E2F1, repressing its expression and therefore inhibiting its effects ^75^. This property has been reported to affect breast cancer tumorigenesis as well. It accompolishes this by transcriptionally activating CCNE1 and CCNE2 leading to G1/S phase transition resulting in increased proliferation ^76^. E2F8 has also been shown to be overexpressed in patient data obtained from TCGA. Patients that had higher expression of E2F8 showed significantly poorer prognosis^77^. Additionally it has been recently shown that post-transcriptional regulation of E2F8 by RNA binding proteins in breast cancer cell lines, increases the proliferative capabilities of breast cancer^78^. It is clear that E2F8 overexpression increases the invasive properties of breast cancer and results in poorer prognosis of breast cancer patients when overexpressed.

BUB1 Mitotic Checkpoint Serine/Threonine Kinase B (BUB1B) has a significant relation with both stage and grade of the tumor. Overexpression of BUB1B is more often noticed in high grade breast cancer and it is correlated with shorter distant metastasis-free survival ^79^. BUB1B’s expression values have been shown to solely divide the breast cancer patients into metastatic and non-metastatic ^80^.

Transmembrane Protein 208 (TMEM208) codes for a transmembrane protein located on endoplasmic reticulum. TMEM208 overexpression has been shown to inhibit ER stress and autophagy^81^. Also, TMEM208 transcription is affected by oxygen levels, especially by hypoxic conditions under which HIF-1α binds to its promoter, resulting in an increase in transcript levels^82^. Hypoxic conditions have been shown to induce invasiveness of breast cancer cells under the orchestration of HIF-1α^83^.

Proteasome 26S Subunit, Non-ATPase 2 (PSMD2) codes for a protein part of the proteasome subunit of the ubiquitin proteasome system (UPS).This gene is upregulated in breast cancer cells as compared with normal adjacent tissue by more than 3 fold^84^. High levels of PSMD2 correlate with shorter over all survival and distant metastasis free survival and the gene is also part of a metastatic signature that relates to poorer prognosis^85,86^. Silencing of PSMD2 inhibits breast cancer cell proliferation, tumor growth and increases apoptosis^85^.

Tubulin Alpha 1b (TUBA1B) is a member of tubulin family proteins, which are major components of microtubules. TUBA1B is involved in a diverse spectrum of biological activities including cell adhesion, movement, replication and division. Although its role is undetermined in breast cancer and metastatic progression, it was shown to be associated with poor survival in liver cancer ^87^.

Ubiquitin Conjugating Enzyme E2 A (UBE2A), also known as RAD6A, is a ubiquitin conjugating enzyme which is involved in protein metabolism. Studies showed that UBE2A is upregulated in aggressive breast cancers and its expression correlates with metastasis ^88^ ^89^. In one study, authors also revealed that increased RAD6 levels are associated with cancer stem cell phenotype in ovarian cancer resulting in disease recurrence and metastasis ^90^.

## Conclusion

Our results suggest that PAM50 plasticity exists in response to treatment. Moreover, metastasized tissue shows different PAM50 class when compared with primary tissue. So, there is a need to reassess the classes in case of relapse and metastasis. For example patients with HER2 enriched phenotype are treated with targeted therapy, but if these tumors switch to a luminal phenotype upon relapse or metastasis, then a second line of treatment can be added to deal with this switch^7^.

Additionally, our suggested metastatic biomarkers can be used to better predict metastatic events in breast cancer patients. Previously in literature several studies have identified metastatic biomarkers in breast cancer^91–93^. But here our markers show prognostic importance in 7 different cohorts.

## Notes

### Competing Interest Statement

The authors have declared no competing interest.

### Summary of Updates

Author name was revised due to a misspelling.

